# *Streptococcus pneumoniae* infection promotes histone H3 dephosphorylation by modulating host PP1 phosphatase

**DOI:** 10.1101/2020.01.14.905968

**Authors:** Wenyang Dong, Orhan Rasid, Christine Chevalier, Michael Connor, Matthew Eldridge, Melanie Anne Hamon

## Abstract

Pathogenic bacteria can alter host gene expression through post-translational modifications of histones. We show for the first time that a natural colonizer, *Streptococcus pneumoniae*, also induces specific histone modifications, including robust dephosphorylation of histone H3 on serine 10, during infection of respiratory epithelial cells. Two bacterial factors are important for the induction of this modification: the bacterial toxin PLY, a pore-forming toxin, and the pyruvate oxidase SpxB, an enzyme responsible for H_2_O_2_ production. The combined effects of PLY and H_2_O_2_ lead to host signaling which culminates in H3S10 dephosphorylation, mediated by the host cell phosphatase PP1. Strikingly, *S. pneumoniae* infection induces dephosphorylation and associated activation of PP1 catalytic activity. Colonization of cells, which lacked active PP1, resulted in the impairment of intracellular *S. pneumoniae* survival. Interestingly, PP1 activation mediating H3S10 dephosphorylation is not restricted to *S. pneumoniae* and appears to be a general epigenomic mechanism favoring intracellular survival.

## Introduction

*Streptococcus pneumoniae* is a Gram-positive, extracellular bacteria that colonizes the human nasopharynx and respiratory tract. It is a leading cause of bacterial pneumonia and meningitis worldwide, particularly in developing nations (1). In the complex relationship with its host, *S. pneumoniae* can act as both an adapted commensal and an invasive pathogen. On the one hand, *S. pneumoniae* may reside asymptomatically in the upper respiratory tract, a phenomenon which is particularly common in children and greatly contributes to transmission. On the other hand, bacteria can breach epithelial barriers, enter the bloodstream, and eventually cause severe invasive pneumococcal diseases (IPDs), such as pneumonia, meningitis and sepsis. To date, 98 serotypes of this encapsulated diplococcus have been described based on variations in their capsule polysaccharide (CPS) (2). However, not all serotypes are equal in their capacity to cause IPDs, only 20 to 30 of them are principally associated with invasiveness diseases, such as serotype 1, 4 and 14 (3,4). In contrast, several serotypes, such as 6B, 11A and 23F, are more likely to be carried for longer periods, and thus regarded as carriage serotypes (3). However, due to the introduction of pneumococcal conjugate vaccines and prominent capsule switching, the serotype prevalence and distribution in IPDs and carriage regularly changes.

For both carriage and invasive pneumococcal strains, the establishment of colonization begins upon contact with host respiratory epithelium. *S. pneumoniae* interacts with epithelial cells via multiple and complex processes, and this multifactorial event has been well characterized with respect the bacterial factors involved. Surface factors, such as CbpA and ChoP, have been reported to participate in adhesion to epithelial cells (2). Another major virulence factor, the pore-forming toxin pneumolysin (PLY), is proposed to be involved in invasion by breaching the epithelium (5). However, the processes involved in pneumococcal-epithelial interaction, such as how *S. pneumoniae* modulates host cell signaling is understudied.

Recent studies show that pathogens reprogram host cells during infection through bacteria-triggered histone modifications, which modulate host transcription (6). In eukaryotic cells, DNA is packaged with histones into chromatin, and the covalent post-translational modifications at the tails of core histones function to dynamically control DNA accessibility, affect the recruitment and stabilization of transcription associated factors, and therefore regulate the cell’s transcriptional programs (7). Bacterial pathogens have been shown to target host histones directly through the secretion of factors targeting chromatin and named nucleomodulins, or indirectly through modulation of host signaling cascades. However, histone modifications induced by a natural colonizer, such as *S. pneumoniae* have not yet been explored.

During bacterial infection, histone H3 was found to be dephosphorylated following the loss of membrane integrity mediated either by the secretion of cholesterol-dependent cytolysin (CDC) toxin or by insertion of translocon from the type III secretion system (8,9). H3S10 dephosphorylation was shown to be induced *in vitro* in epithelial cells, and to correlate with the repression of inflammatory genes. Although cellular processes, such as entry into mitosis or activation by extracellular signals (EGF, cellular stress, or inflammation) have been shown to associate with H3S10 phosphorylation, its role is unclear (10-12). Key kinase signaling pathways, including MAPK and NF-κB, induce the phosphorylation of histone H3 on residue 10 (H3S10) or 28 (H3S28), which are important for crosstalk with histone H3 acetylation, a canonical mark of transcriptional activation (13,14). In contrast, how H3S10 becomes dephosphorylated is less understood. Furthermore, the molecular basis and the role of this histone modification during bacterial infection remain unknown.

In this manuscript, we show for the first time that *S. pneumoniae* induces histone modifications in host lung epithelial cells, both *in vitro* and *in vivo*. We identify the CDC toxin PLY as a key factor in mediating H3 dephosphorylation, but also reveal the strong contribution of the pyruvate oxidase, SpxB. Both factors PLY and SpxB activate the host phosphatase PP1, which we show is responsible for H3 dephosphorylation. Interestingly, we find that infection triggers dephosphorylation of PP1 on threonine 320 (T320) which is required for its activation, and that this process is necessary to permit efficient intracellular infection. By describing the molecular basis and the role of H3S10 dephosphorylation, we illustrate a common mechanism of host alteration by bacteria relevant during both colonization and infection.

## Results

### *Streptococcus pneumoniae* induces dephosphorylation of histone H3 on serine 10 during infection

Histone H3 dephosphorylation on serine 10 was initially observed following treatment of epithelial cells with purified bacterial toxins, including PLY of *S. pneumoniae* (8). In order to evaluate whether histone H3 phosphorylation is modulated by bacteria during infection, we exposed A549 lung epithelial cells to two different serotypes of *S. pneumoniae* (R6 or TIGR4) at increasing multiplicities of infection (MOI) and assessed the phosphorylation of H3S10 by immunoblotting. As a control, total levels of H3 and actin were also measured. Data in Figure 1A shows that H3S10 is strongly dephosphorylated, and this effect is proportional to the number of bacteria used in the infection. We further validated that this effect was not only restricted to alveolar epithelial A549 cells, as bronchial epithelial Beas-2B cells displayed the same reduction in H3S10ph levels (Figure S1A). H3T3, H3T11 and H3S28 were also dephosphorylated in response to bacteria, however H3T6 was not, suggesting only specific residues are targeted (Figure S1B). Since H3S10 phosphorylation has been previously associated with bacterial infection (8,15,16), we focused our study on this modification.

**Fig.1.**
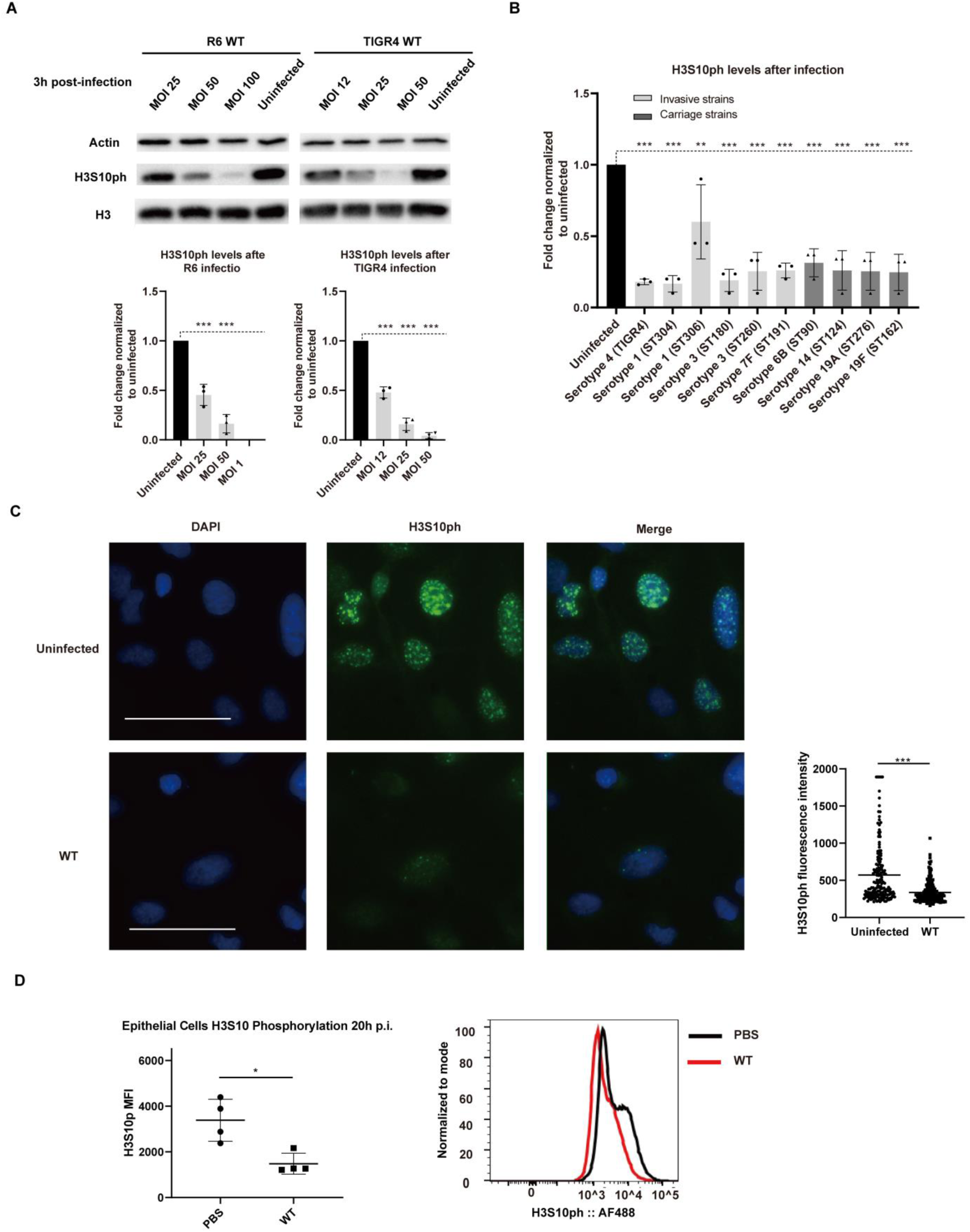
*Streptococcus pneumoniae* induces specific dephosphorylation of histone H3 on serine 10 independently of the cell cycle. **(A)** Phosphorylation levels of histone H3S10 as detected by immunoblotting in uninfected A549 cells and cells infected with *S. pneumonia* strain R6 or TIGR4 at the indicated multiplicity of infection (MOI). Quantification of H3S10ph immunoblots, normalized first to total actin levels and then to the uninfected sample. Error bars represent SD of 3 independent experiments. Statistical significance was calculated using One-way ANOVA method (Dunnett’s Post Hoc test, uninfected as control group), ***P < 0.001. **(B)** Quantification of levels of H3S10 phosphorylation detected by immunoblotting in uninfected A549 cells and cells infected with TIGR4 or clinical *S. pneumonia* strains (from carriage or invasiveness), serotypes and MLST types at MOI 25 are indicated. Error bars represent SD of 3 independent experiments. Statistical significance was calculated using One-way ANOVA method (Dunnett’s Post Hoc test, uninfected as control group), **P < 0.01, ***P < 0.001. **(C)** H3S10ph was detected by immunofluorescence in Beas-2B cells under uninfected and 1h TIGR4 (MOI=25) infected conditions. Size bars represent 50 µm. On the right, the quantification of nuclear fluorescence intensity integrates independent duplicate experiments (n ≥ 200 cells per condition). Statistical significance was calculated using a Student’s t test, ***P < 0.001. **(D)** H3S10ph levels of lung epithelial cells from mice treated with PBS or infected with *S. pneumonia* TIGR4 (5×10^6 CFU) for 20 hours were analyzed by FACS. Mean fluorescence intensity (MFI) is represented with error bars indicating SD, n = 4 mice per condition. Statistical significance was calculated using a Student’s t test, *P < 0.05.

Given that H3S10 dephosphorylation was observed during infection with two different laboratory strains, we hypothesized this effect was a shared mechanism among *S. pneumonia* serotypes. We assessed clonal representatives of different invasive and carriage serotypes for their ability to induce H3S10 dephosphorylation. We show that all serotypes tested induced strong H3S10 dephosphorylation in A549 cells (Figure 1B). These results demonstrate that reduction of H3S10ph levels is a common feature of *S. pneumoniae* infection and is induced by clinical isolates as well as laboratory strains.

H3S10ph has been associated with the mitotic phase of the cell cycle (17), therefore we investigated whether the observed loss reflected changes in the cell cycle induced by infection. Cells were synchronized using thymidine, infected upon release, and the proportion of cells in each stage of the cell cycle was measured using propidium iodine staining. FACS analysis of stained cells showed no significant difference in the percentage of cells in each stage between uninfected and infected cells (Figure S1C), strongly suggesting that infection does not affect the cell cycle. Furthermore, immunofluorescence experiments were performed to determine H3S10ph levels in non-mitotic cells. Nuclear staining with DAPI clearly distinguishes cell in interphase versus mitosis and allows direct visualization of interphase cells, this confirmed that many cells, which do not have condensed chromosomes, stain positive for H3S10ph (Figure 1C). Quantification of fluorescence intensity upon infection with *S. pneumoniae* shows a significant reduction compared to uninfected cells (Figure 1C). These data further support the finding that infection-induced dephosphorylation occurs in interphase cells, independently of the cell cycle.

We next determined whether H3S10 dephosphorylation occurred during *in vivo* infection. We performed intranasal inoculation of mice with *S. pneumoniae* and used antibodies against cell lineage markers to specific cell types isolated from collected mouse lungs (Figure S1D). The levels of H3S10 phosphorylation in epithelial cells after 24h of infection were further evaluated by FACS analysis. In comparing infected to uninfected cells, a shift in the fluorescence level of a large proportion of epithelial cells was observed (Figure 1D). Interestingly, dephosphorylation was mainly detected in epithelial cells, and other monitored cells did not display significant changes in this histone mark (Figure S1E). These data further support that H3 dephosphorylation occurs in terminally differentiated cells, independently of the cell cycle. Therefore *S. pneumoniae* induces specific H3S10 dephosphorylation in epithelial cells during infection.

### *S. pneumoniae* toxin PLY is important for H3S10 dephosphorylation

The cholesterol dependent cytolysin of *S. pneumoniae*, PLY, was previously shown to induce H3S10 dephosphorylation when treating epithelial cells with purified toxin (8). We thus hypothesized that PLY was the main factor responsible for this modification during infection. Chromosomal deletion mutants of PLY were generated in both R6 and TIGR4 strains of *S. pneumoniae* and tested for their ability to dephosphorylate H3. Immunoblotting experiments show that H3 dephosphorylation is only partially blocked with the Δ*ply* mutant as compared with wild type infection (Figure 2A). Since PLY does not have a signal sequence for secretion, it is thought that the toxin is release upon lysis of bacteria in a manner dependent on the autolysin LytA. We therefore generated a *lytA* deletion mutant, which releases less PLY than wild type bacteria, and induces a similar dephosphorylation of H3 as a Δ*ply* mutant (Figure S2A, B). Therefore, H3 dephosphorylation is in part mediated by PLY, which is released upon LytA dependent lysis of a subpopulation of bacteria.

**Fig.2.**
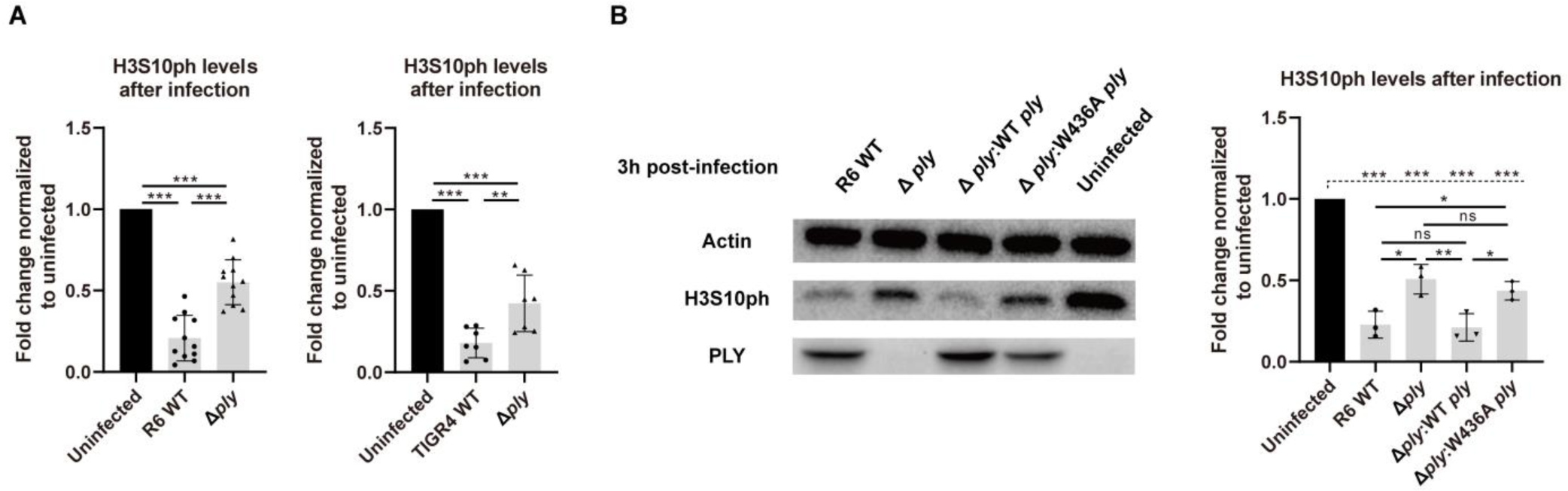
*S. pneumoniae* toxin PLY is important for H3S10 dephosphorylation. **(A)** Quantification of H3S10ph in uninfected cells, cells infected with wild-type *S. pneumonia* strains R6 (MOI=50) and TIGR4 (MOI=25), and their respective *Δply* mutant strains for 3 h. **(B)** Representative immunoblots of cells infected with wild-type *S. pneumonia* (R6 strain), a *Δply* mutant, a *Δply* mutant complemented with wild-type *ply*, or a *ply* point mutant without pore-forming activity (W436A) at MOI 50 for 3 h. All quantifications in graphs show the mean +/− SD of at least 3 independent experiments. Statistical significance was calculated using One-way ANOVA method (Turkey Post Hoc test when compared within infection groups as indicated with solid line, or Dunnett’s Post Hoc test when compared to uninfected control as indicated with dotted line), *P < 0.05, **P < 0.01, ***P < 0.001.

To understand how PLY was inducing histone modification, we complemented the Δ*ply* mutant with a PLY bearing a point mutation, W436A, rendering it non-hemolytic (18). We compared the level of H3S10 phosphorylation of this strain to that of a strain complemented with wild type PLY. The wild type complement induced the same level of H3S10 dephosphorylation as wild type bacteria, whereas the W436A mutant phenocopied the Δ*ply* mutant (Figure 2B). Therefore, PLY contributes to dephosphorylation of H3 in a pore formation-dependent manner. Given that the level of H3S10ph was clearly not restored to uninfected levels upon infection with the Δ*ply* mutant, other additional contributing bacterial factors are probably also responsible for inducing this histone modification.

### H_2_O_2_ generated by SpxB is required for *S. pneumoniae* mediated H3S10 dephosphorylation

Reports have shown that stress response, such as oxidative stress, can lead to H3 dephosphorylation (19,20). Interestingly, *S. pneumoniae* expresses a pyruvate oxidase that converts pyruvate to acetyl-phosphate and generates H_2_O_2_ as a byproduct. The produced H_2_O_2_, which is a molecule freely diffusible across membranes, is high enough to detect in the supernatant of infected cells, and a deletion of the SpxB oxidase results in a complete block in H_2_O_2_ production (Figure S3A). To determine whether SpxB was contributing to H3 dephosphorylation, we infected cells with a deletion mutant. In both strains R6 and Tigr4 H3 dephosphorylation was prevented upon infection with a Δ*spxB* mutant. In fact, H3 levels were comparable to the uninfected level upon infection with Δ*spxB* or a double Δ*ply*Δ*spxB* mutant (Figure 3A). Additionally we performed immunofluorescence experiments with the double Δ*ply*Δ*spxB* mutant, and quantification of H3S10ph fluorescence intensity shows that the double mutant does not induce dephosphorylation (Figure 3B).

**Fig.3.**
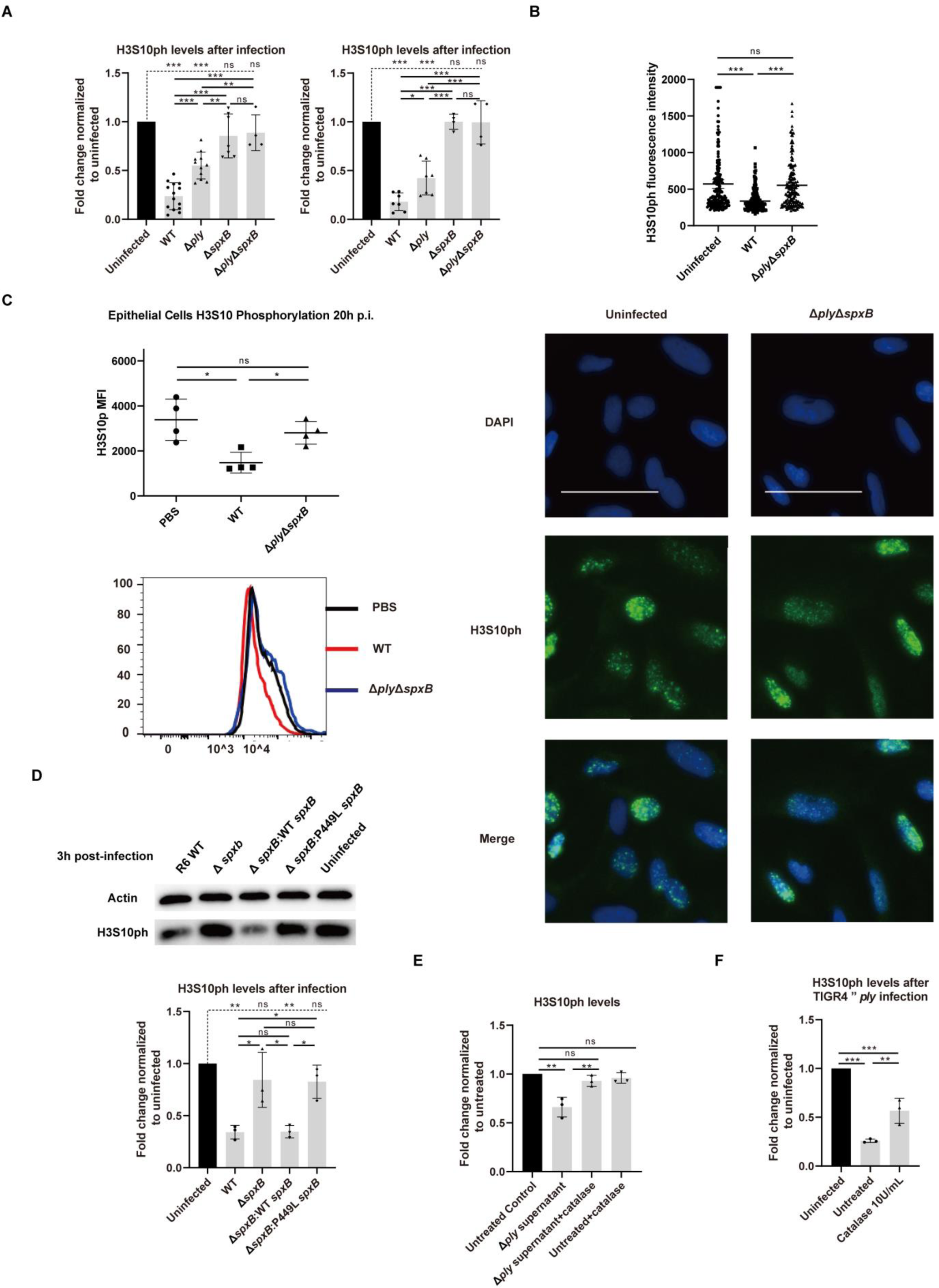
H2O2 generated by SpxB is required for *S. pneumoniae* mediated H3S10 dephosphorylation. **(A)** Quantification of H3S10ph in uninfected cells, cells infected with wild-type *S. pneumonia* strains R6 (MOI=50) and TIGR4 (MOI=25), a *Δply* mutant, a *ΔspxB* mutant, or a *ΔplyΔspxB* double mutant for 3 h. **(B)** H3S10ph was detected by immunofluorescence in Beas-2B cells under uninfected and 1h (MOI=25) infected conditions. On the right, the quantification of nuclear fluorescence intensity integrates independent duplicate experiments (n ≥ 200 cells per condition). Size bars represent 50 µm. **(C)** H3S10ph levels of lung epithelial cells from mice treated with PBS,wild-type *S. pneumonia* TIGR4 (5×10^6 CFU), or a *ΔplyΔspxB* double mutant (5×10^6 CFU) for 20 hours were analyzed by FACS. Mean fluorescence intensity (MFI) is represented with error bars indicating SD, n = 4 mice per condition. **(D)** Immunoblot images are shown on the left and quantifications on the right. The left image is shown for representative immunoblots of cells infected with wild-type *S. pneumonia* (R6 strain), a *ΔspxB* mutant, a *ΔspxB* mutant complemented with wild-type *spxB*, or a *spxB* point mutant without catalytic activity (P449L) at MOI 50 for 3 h. **(E)** Cells were incubated 2h with the filter-sterilized supernatant from infection wells, or with 15 min catalase pretreated filter-sterilized supernatant, or 15 min catalase pretreated cell culture medium, respectively. **(F)** Quantification of H3S10ph in uninfected cells, cells infected with TIGR4 *ply* mutant (with or without catalase during infection) at MOI 25 for 3 h. Error bars in the quantifications represent SD of at least 3 independent experiments. Statistical significance was calculated using One-way ANOVA method (Turkey Post Hoc test when compared within infection groups as indicated with solid line, or Dunnett’s Post Hoc test when compared to uninfected control as indicated with dotted line), *P < 0.05, **P < 0.01, ***P < 0.001.

To determine whether *S. pneumoniae* mediated H3 dephosphorylation requires the catalytic activity of SpxB and H_2_O_2_, the byproduct of catalysis, we generated a Δ*spxB* mutant complemented with wild type *spxB* and or a catalytically inactive *spxB* (*spxB* P449L) (21). Both strains were used in infection and the levels of H3S10ph were determined by immunoblotting. Whereas Δ*spxB:spxB* was able to restore H3 dephosphorylation to wild type levels, the catalytically inactive complement, Δ*spxB:spxBP449L* did not (Figure 3D). To further show that it is the byproduct of SpxB catalysis, H_2_O_2_, rather than other metabolic intermediates, which is important for H3 dephosphorylation, we performed experiments in the presence of catalase. Catalase is an enzyme that converts H_2_O_2_ to water and oxygen and thereby neutralizing the downstream effects H_2_O_2_. Cells infected with Δ*ply* TIGR4 or treated with infection supernatant displayed significant H3 dephosphorylation (Figure 3E and 3F). However, upon catalase treatment H3 dephosphorylation was significantly impaired, reinforcing the finding that H_2_O_2_ produced by SpxB is the main driver for H3 dephosphorylation. In fact, H_2_O_2_ alone induces H3 dephosphorylation in A549 cells in a dose dependent manner (Figure S3B). Therefore H_2_O_2_ generated by SpxB is a factor contributing to H3 dephosphorylation in addition to the pore-forming toxin PLY.

In vivo experiments were performed to determine whether SpxB and PLY were necessary for H3 dephosphorylation in lung epithelial cells. Whereas wild type infection leads to a significant shift in phosphorylation H3 levels, the double Δ*ply*Δ*spxB* mutant does not. In fact, H3S10 dephosphorylation did not occur when inoculating mice intranasally with the Δ*ply*Δ*spxB* mutant (Figure 3C). Therefore both PLY and SpxB are required to induce H3 dephosphorylation upon infection both *in vitro* and *in vivo* in epithelial cells. We noticed that a single *ΔspxB* mutant does not induce greater H3S10 dephosphorylation than a double Δ*ply*Δ*spxB* mutant. This could be explained by the finding that a mutant in SpxB is diminished in PLY release and pore-formation on epithelial cells (22).

### PP1 is the host phosphatase mediating H3S10 dephosphorylation

Given that neither PLY nor SpxB are directly targeting H3S10 for modification, we searched for a host phosphatase which could mediate the observed dephosphorylation. We first used chemical inhibitors which target known phosphatases; okadaic acid a potent and selective inhibitor of protein phosphatase 1 (PP1) and 2A (PP2A), with a greater potency for 2A, and tautomycetin which specifically inhibits PP1. Cells were pretreated with the two inhibitors and subsequently infected with either R6 or TIGR4 strains of *S. pneumoniae*. Detection of phosphorylated H3 levels by immunoblotting show that okadaic acid had no effect, whereas tautomycetin blocked infection-induced dephosphorylation (Figure 4A and S4A). These results were further validated by RNA interference using siRNA targeting all isoforms of PP1 (α, β, γ). The knock down efficiency siRNA clearly shows that the level of all isoforms is equally decreased (Figure S4B). Under these conditions H3S10 dephosphorylation was fully blocked upon infection, similar to the results obtained with tautomycetin (Figure 4B). Consistent with PP1 being the main phosphatase targeting H3, siRNA of PP1 also fully blocked H_2_O_2_ and purified PLY mediated H3 dephosphorylation (Figure 4B, 4C, 4D). Together these results show that PP1 is the phosphatase involved in dephosphorylating H3 and that both PLY and SpxB mediate this effect through the same host pathway.

**Fig.4.**
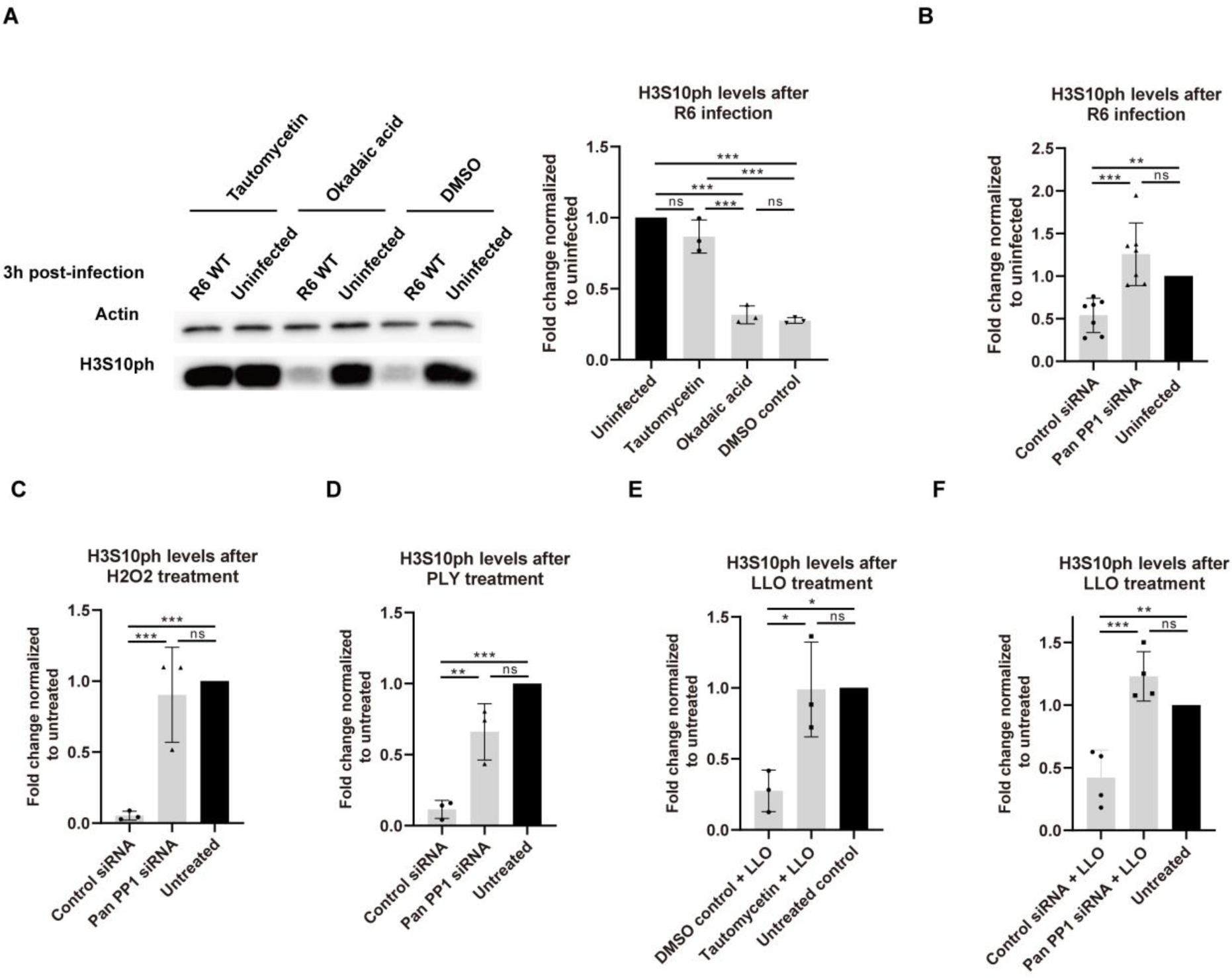
PP1 is the host phosphatase mediating H3S10 dephosphorylation. **(A)** Representative immunoblots images are shown on the left and quantifications on the right. A549 cells pretreated with PP1 inhibitor Tautomycetin, PP2A inhibitor Okadaic acid or DMSO control and infected with R6 (MOI=50) for 3h. **(B)** A549 cells are transfected with PP1 siRNA or control siRNA prior to 3h R6 (MOI=50) infection. **(C)** A549 cells are transfected with PP1 siRNA or control siRNA prior to PLY treatment for 30 min. **(D)** A549 cells are transfected with PP1 siRNA or control siRNA prior to H2O2 treatment for 1h. **(E)** Hela cells are pretreated with PP1 inhibitor Tautomycetin prior to LLO treatment for 20 min. **(f)** Hela cells are transfected with PP1 siRNA or control siRNA prior to LLO treatment for 20 min. All quantification graphs above are of at least 3 independent experiments. All quantifications in graphs show the mean +/− SD and statistical significance was calculated using One-way ANOVA method (Turkey Post Hoc test), **P < 0.01, ***P < 0.001.

Previous studies have demonstrated that other toxins of the CDC family, such as listeriolysin O (LLO) from *Listeria monocytogenes*, also induced dephosphorylation of H3S10, but had not identified the mechanism at play (8). Therefore we tested whether PP1 was also required for LLO induced H3 dephosphorylation. Cells were treated with purified LLO and H3 dephosphorylation was observed in HeLa cells (Figure 4E and 4F). However, upon pretreating cells with either tautomycetin or by silencing the expression PP1 with siRNA, H3 dephosphorylation was blocked (Figure 4E and 4F). Therefore, PP1 is modulated by at least two bacterial toxins mediating H3S10 dephosphorylation.

### Bacterial infection induces dephosphorylation of PP1

In its resting state PP1 is phosphorylated on T320 and dephosphorylation of this residue correlates with PP1 activation. Therefore we determined whether PP1 was being activated by infection through dephosphorylation of T320. Firstly, lysates from infected and non-infected cells were probed by immunoblotting for the total levels of each PP1 isoform (Figure 5A). Importantly, the total level of PP1 is unaltered by infection regardless of the mutant used. In contrast, the level of phosphorylated PP1 is significantly decreased upon infection with WT *S. pneumoniae*. Interestingly, PP1 dephosphorylation was partially rescued upon infection with a Δ*ply* mutant, and fully prevented upon infection with a Δ*spxB* or the double Δ*ply*Δ*spxB* mutant. Therefore the levels of phosphorylated PP1 fully correlated with the levels of phosphorylated H3S10 (Figure 5B), strongly suggesting that activation of PP1 by dephosphorylation was leading to dephosphorylation of H3.

**Fig. 5.**
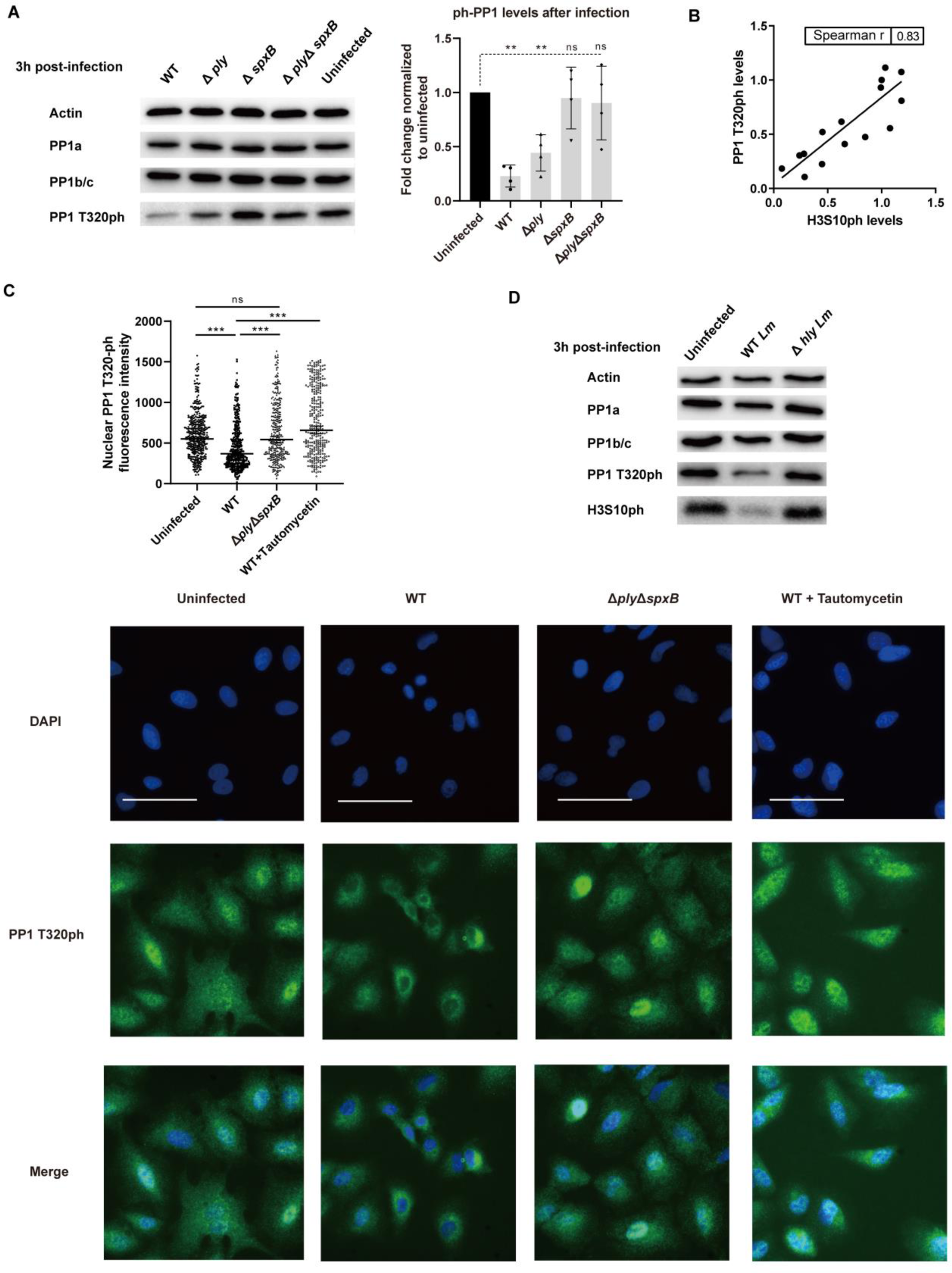
Bacterial infection induces dephosphorylation of PP1. **(A)** A549 cells were infected for 3 h with wild-type and indicated mutants of *S. pneumonia* strain TIGR4 (MOI=25). A representative immunoblot (left) and a quantification (right) of 4 independent experiments are shown. The PP1 T320ph levels are normalized to PP1a and to the uninfected control condition. *S. pneumonia*. Error bars represent SD and statistical significance was calculated using One-way ANOVA method (Dunnett’s Post Hoc test, uninfected as control group), **P < 0.01. **(B)** The correlation of H3S10ph levels and PP1 T320ph levels of infected cells from 4 independent experiments is calculated using nonparametric Spearman’s correlation coefficient method **(C)** Immunofluorescence of PP1 T320ph in A549 cells under uninfected, 3h wild-type TIGR4 infection, 3h *ΔplyΔspxB* double mutant infection and 3h wild -type infection at MOI 25 with 3h Tautomycetin pretreatment conditions. Size bars represent 50 µm. Quantification of nuclear PP1 T320ph fluorescence intensity from at least 2 independent experiments, for more than 300 cells counted each condition. (**D**) HeLa cells were infected for 3 h with the wild-type *Listeria* (EGD strain) and its *hly* mutant at MOI 50. A representative immunoblot of 3 independent experiments is shown.

We confirmed this finding by immunofluorescence by evaluating the level of phosphorylated PP1 in A549 cells. Interestingly, PP1 became dephosphorylated specifically in the nucleus of cells infected with WT bacteria, where histone H3 dephosphorylation occurs (Figure 5C). In contrast, the levels of nuclear phosphorylated PP1 did not change upon infection with the double Δ*ply*Δ*spxB* mutant. Since tautomycetin blocked infection-induced H3 dephosphorylation, we measured the corresponding levels of phosphorylated PP1. Tautomycetin fully blocked infection-induced PP1 dephosphorylation, and appeared to induce phosphorylation (Figure 5C, Figure S5A). These results show that the effect of PLY and SpxB converge on PP1, which likely auto-dephosphorylates itself.

Our data show that other toxins in the CDC family, such as LLO, mediate H3 dephosphorylation through PP1. We therefore wanted to determine whether infection with *Listeria monocytogenes*, which produces LLO, also induced PP1 T320 dephosphorylation. To address this point, we infected HeLa cells with either wild type *L. monocytogenes*, or a mutant lacking the LLO toxin (Δ*hly*). The representative immunoblots in Figure 5D show that WT bacteria induces dephosphorylation of both H3 and PP1. However, infection with the Δ*hly* strain did not induce dephosphorylation. Therefore dephosphorylation of PP1, leading to H3 dephosphorylation, is a common mechanism activated by bacteria that produce cholesterol dependent cytolysins, and is essential for H3 dephosphorylation.

Surprisingly, although infection activates PP1, which is a pleotropic enzyme, the general phosphorylation levels of the host do not seem to be altered. Indeed, we observed no significant change in the total levels of phosphorylated serine and threonine in total cell lysates upon infection (Figure S5B). Furthermore, we tested if activated PP1 dephosphorylated non-histone substrates, such as AKT, a kinase known to be regulated by PP1 (23,24). AKT phosphorylation at S473 was not altered during *S. pneumoniae* infection (Figure S5B). Therefore, PP1 activation by infection seems to have some degree of substrate specificity. However, we cannot rule out that other proteins besides histone H3 are dephosphorylated during infection.

### H3S10 dephosphorylation correlates with transcriptional repression of inflammatory genes, but is not required

Beyond a correlation with the cell cycle, H3 phosphorylation has been associated with transcriptional activation of inflammatory genes downstream of LPS stimulation (25). We therefore hypothesized that the role of H3 dephosphorylation could be to downregulate inflammatory genes during infection. To test this, we tested the expression of a panel of 26 pro-inflammatory genes following infection with wild type *S. pneumoniae*, which induces H3 dephosphorylation, and a double Δ*ply*Δ*spxB* mutant, which does not. We focused on genes that were differentially regulated between WT and the Δ*ply*Δ*spxB* mutant, and found seven: *CCL2, CCL4, CCL5, CXCL1, TNF, CSF2*, and *IL1A*. Strikingly, while infection with wild type did not significantly change the level of gene expression, the double mutant increased gene expression compared to uninfected cells (Figure 6). In contrast expression levels of a control gene, HPRT1, were not altered. These results suggest that *S. pneumoniae* actively suppressed inflammatory gene transcription in epithelial cells, in a manner that is dependent on PLY and SpxB.

**Fig.6.**
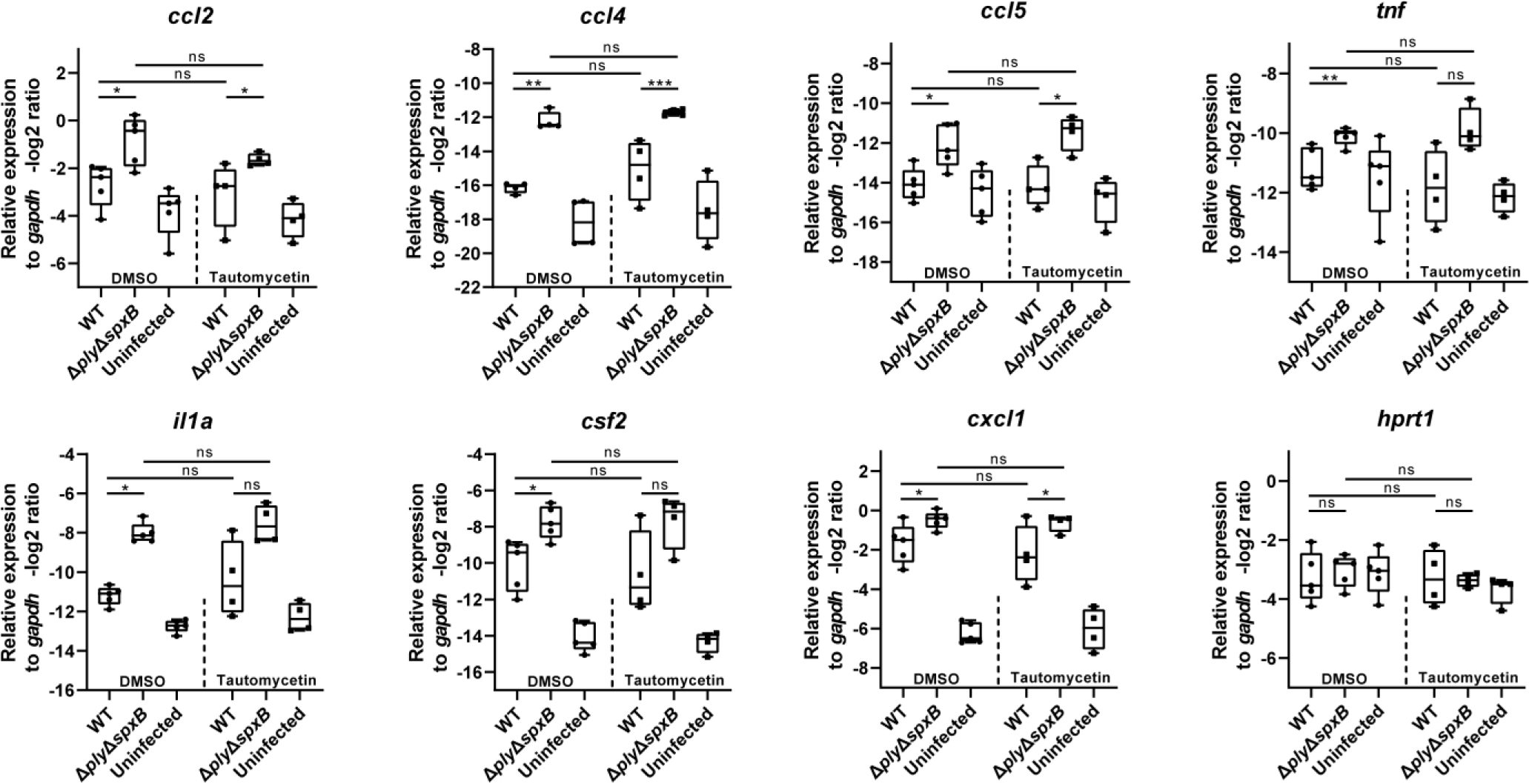
H3S10 dephosphorylation correlates with transcriptional repression of inflammatory genes, but is not required. Untreated cells or Tautomycetin pretreated cells are infected by wild-type TIGR4 *S. pneumonia* and its *ΔplyΔspxB* double mutant at MOI 25. RNA was extracted and qRT–PCR was performed to analyze the relative expression of indicated inflammatory genes and housekeeping gene *hprt1*. The gene expression levels are shown as -log2 relative to control gene *gapdh*, the results are shown as mean and SD of 4 independent experiments. For statistical calculation, -log2 form data is first converted to the linear form by the 2^ (-ΔCT) calculation, and then tested with a Student’s t test method, *P < 0.05, **P < 0.01, ***P < 0.001.

To determine whether *S. pneumoniae* mediated gene suppression occurred through H3 dephosphorylation, we repeated RT-PCR analyses in the presence of the PP1 inhibitor tautomycetin. Indeed, since tautomycetin fully blocks H3 dephosphorylation, differential gene expression between WT and the Δ*ply*Δ*spxB* mutant would also be blocked in the presence of this inhibitor. However, treatment with tautomycetin did not change the transcriptional repression observed upon infection with wild type bacteria for the genes tested (Figure 6). These results suggest that although H3 dephosphorylation is correlated with the repression of inflammatory genes, it is not required for it. We cannot exclude that H3 dephosphorylation could be mediating transcriptional repression of other genes, which we have not yet identified.

### Blocking H3S10 dephosphorylation through PP1 inhibition impairs efficient intracellular infection

To determine whether PP1-mediated dephosphorylation was necessary for bacterial infection, we assessed the impact of blocking PP1 catalytic activity, using the chemical inhibitor tautomycetin, on infection. *In vitro S. pneumoniae* mainly remains adhered to the outside of epithelial cells (26,27). In some instances though, it is able to invade and survive a short period of time (approximately 24h) inside epithelial cells, a mechanism that could be important for the “microinvasion” observed *in vivo* (5,28). We first measured the effect of PP1 on bacterial adherence. We compared the number of bacteria recovered from cells treated or not with tautomycetin, the PP1 inhibitor. Results in Figure 7A show that there is no significant difference between the two conditions. Similar results were obtained in evaluating the number of Δ*ply*Δ*spxB* mutant bacteria in the presence or absence of tautomycetin (Figure 7A). We also evaluated the number of intracellular bacteria at 4 and 6 hours post infection. Strikingly, upon inhibition of PP1 a slight, but significant decrease in the number of recovered intracellular bacteria was observed at both time points (Figure 7B). These results would suggest that PP1 is important for intracellular bacterial survival. We further tested the effect of PP1 inhibition on infection with the double Δ*ply*Δ*spxB* mutant. This mutant does not modify PP1 and no H3 dephosphorylation occurs. Interestingly, tautomycetin has no effect on intracellular survival of Δ*ply*Δ*spxB* mutant bacteria indicating that this inhibitor is specifically targeting PP1 activity (Figure 7B).

**Fig.7.**
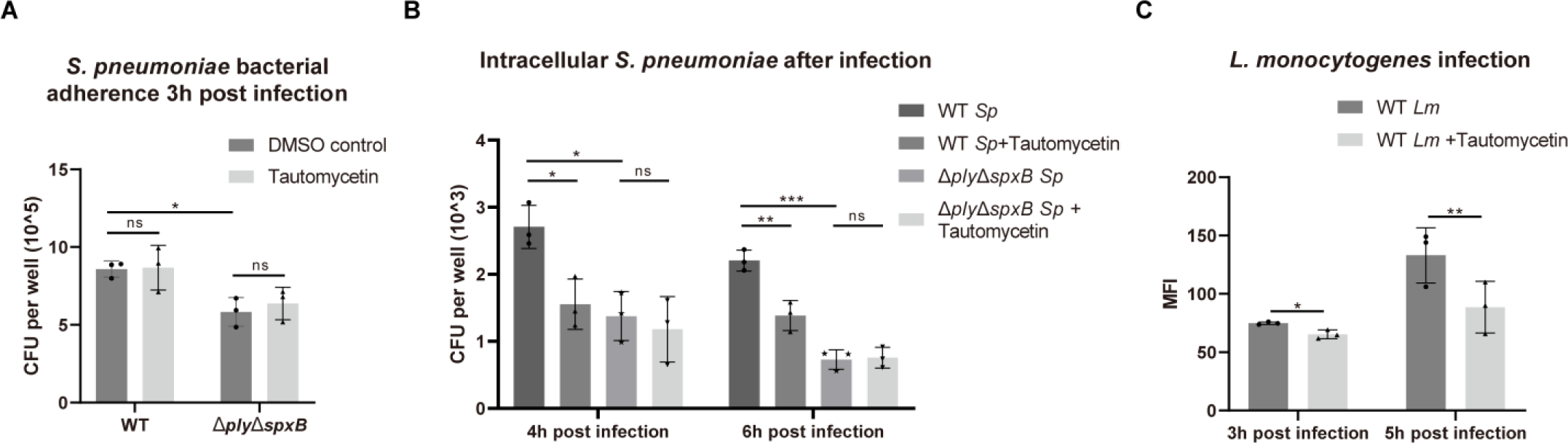
H3S10 dephosphorylation is necessary for efficient bacterial infection. Wild-type *S. pneumonia* (R6) and its *ΔplyΔspxB* double mutant were incubated with A549 epithelial cells at MOI 50 and assayed for either bacterial adherence **(A)** or intracellular bacteria **(B)**. The results are mean +/− SD from 3 independent experiments and statistical significance was calculated using Student’s t test method, *P < 0.05. **(c)** Caco2 cells were infected with *L. monocytogenes* expressing GFP for the indicated times. Intracellular bacteria were detected by FACS analysis, and the mean fluorescence of intensity (MFI) was calculated. Results are mean +/− SD over 3 independent experiments. In each experiment the ratio between inhibitor group and untreated control group was calculated and the statistical significance was analyzed using Student’s t test method, *P < 0.05, **P < 0.01.

Since *L. monocytogenes*, a facultative intracellular pathogen, also activates PP1 to dephosphorylate H3, we assessed the impact of PP1 inhibition on its intracellular survival. Cells were infected with *L. monocytogenes* expressing GFP and fluorescence intensity of infected cells was determined by FACS analysis. Interestingly, tautomycetin treated cells were less infected than non-treated cells further supporting a role for PP1 in intracellular bacterial survival (Figure 7C). Therefore, bacteria triggered PP1 activation, and probably the downstream H3 dephosphorylation, contribute to optimal intracellular infection of at least two unrelated bacteria.

## Discussion

We report here that histone H3 dephosphorylation is a common modification induced not only by pathogenic bacteria, but also colonizing pneumococci. In fact the bacterial factors involved in inducing this modification, PLY and SpxB, are common to all *S. pneumoniae* serotypes. Importantly, we demonstrate with *in vivo* samples that epithelial cells, which are the first line of response to bacteria entering an organism, are the main cell type affected by this modification. Furthermore, the ability of bacteria to induce active PP1 and thereby H3 dephosphorylation provides an advantage and is necessary for a productive infection.

The role of epithelial cells in a pneumococcal infection has often been overshadowed by the study of “professional” immune cells. However, more than just barrier cells, epithelial cells play a pivotal role in dictating pulmonary innate immune responses upon infection (29). In fact, selectively inhibiting canonical pathways such as NF-kB signaling in epithelial cells results in an impairment in neutrophil recruitment and poor activation of innate immune responses (30). Furthermore, the epithelial barrier is responsible for the production of antibacterial components such as defensins, chemokines and toxic metabolic byproducts (29). Very recently, *S. pneumoniae* at mucosal surfaces was shown to shape epithelial transcriptomic response including innate signaling and regulatory pathways, inflammatory mediators, cellular metabolism and stress response genes (27). Since histone modifications play an important role in regulating gene expression, their modification by bacteria has the potential to significantly alter host responses. A careful study of epithelial cell subtypes would define whether H3 dephosphorylation is found in epithelial progenitor cells or only in specialized and terminally differentiated cells. Such studies would be important to determine whether cells experiencing H3 dephosphorylation are turned over or maintained beyond the presence of bacteria. Our findings that pneumococci modify host histones through active post-translational modification of a host phosphatase, which gives a growth advantage to bacteria, have far reaching implications for both virulent and colonizing serotypes.

Cell intoxication by cholesterol dependent cytolysins (CDC), including PLY, results in characteristic responses which are hallmarks of membrane damage, including ion fluxes, loss of cytoplasm, and host cell death (31). However, at sublethal doses, membrane repair and cell signaling are also important features. Under the conditions used in this study, we do not observe a significant amount of cytotoxicity to host cells, as determined by normal cellular morphology (Fig 1) and unaltered cell cycle upon infection (Fig S1). In agreement with these findings, *in vivo* infections with TIGR4 do not induce noticeable epithelial barrier damage and influx of inflammatory cells. Therefore, H3 dephosphorylation is most likely not linked to *S. pneumoniae* induced cell death.

H3S10 phosphorylation is the most common histone modification linked with bacterial infection. The activation of both NF-κB and MAPKs, which are the major host cell response pathways to infection, lead to H3S10 phosphorylation and associated histone acetylation, resulting in transcriptional activation of target genes (13,14). Several bacterial pathogens have developed strategies to prevent H3S10 phosphorylation by disrupting pro-inflammatory kinase signaling pathways, for example, lethal toxin (LT) from *Bacillus anthracis* and type III secretion system effector OspF from *Shigella flexneri*. Both inactivate MAPKs during infection, subsequently abrogating histone H3S10 phosphorylation, which correlates with repressing inflammatory genes (16,32). However bacteria producing CDC toxins are the only ones to induce dephosphorylation, rather than a block in phosphorylation, which implies that the basal levels of modified histones are changed. In addition to CDC toxins, we show that production of H_2_O_2_ by *S. pneumoniae* amplifies the observed dephosphorylation. Interestingly, SpxB was shown to affect the level of PLY release and epithelial cell pore-formation, indicating that the two proteins share a close link, even though the mechanism is unknown (33). These observations could explain the phenotype of the *ΔspxB* mutant, which displays levels of H3 phosphorylation similar to the double *ΔspxBΔply* mutant and to uninfected cells. The production of H_2_O_2_, a freely diffusible molecule, as metabolic byproduct has been reported for several other bacterial species, including both opportunistic pathogens, such as streptococci, and probiotic or commensal enterococci and lactobacilli (34). Therefore, all H_2_O_2_ producing bacteria have the potential to induce H3 dephosphorylation and alter host cell responses.

Previous attempts to link H3 dephosphorylation with a cellular effect resulted in correlative studies. We previously correlated this modification with a loss of expression of a subset of host genes (8). However, to establish a cause and effect link, inhibiting H3 dephosphorylation is necessary and only possible through inhibition of the enzyme responsible for the modification. We report here that the chemical inhibitor tautomycetin blocks H3 dephosphorylation, while previous attempts to identify such an inhibitor had failed (8). This important step allows us to determine that during infection with *S. pneumoniae*, PP1 mediated H3S10 dephosphorylation is not necessary for the difference in expression of inflammatory genes between wild type and mutant strains. Since H3 phosphorylation is tightly linked to transcriptional regulation (10-12), we hypothesize that *S. pneumoniae* infection may alter the expression of unidentified genes controlled by basal H3S10 phosphorylation in epithelial cells. Further studies would be necessary to identify the genes involved.

The protein phosphatase PP1 is a major protein serine/threonine phosphatases in eukaryotic cells (35). PP1 has been shown to target phosphorylated H3S10 mainly after mitosis, when H3S10 become dephosphorylated (36). The activity of PP1 is regulated through phosphorylation of T320, which is an inhibitory modification probably masking the PP1 active site, and blocking PP1 substrates accessibility (37). A previous report suggested that ultraviolet irradiation induced dephosphorylation of PP1, leading to its activation and triggering subsequent histone H3T11 dephosphorylation (20). Our data highlights that bacterial infection is able to activate PP1 through dephosphorylation, a mechanism used for its own benefit. Notably, the pathways regulated by the PP1-H3 dephosphorylation axis are conserved between at least two Gram positive bacteria, *L. monocytogenes* and *S. pneumoniae*, and are required for efficient intracellular infection in both models. Given the pleotropic role of PP1, we cannot fully rule out that inhibition of PP1 with tautomycetin does not block dephosphorylation of other factors besides histone H3. This would imply that the impairment of intracellular infection could involve other hosts signaling pathways. However, our data show that general dephosphorylation upon infection does not occur and that tautomycetin-mediated impairment of infection only occurs with wild type bacteria, not mutant strains. Although the impact of PP1-H3 dephosphorylation on bacterial infection is slight, our findings do validate a concept that targeting host chromatin modifying enzymes alters bacterial infection. Therefore, our study characterizes and extends the knowledge on a conserved mechanism of host subversion aimed at downplaying host response to invading bacteria.

## Acknowledgments

Work in the M.A.H. laboratory received financial support from Institut Pasteur and the National Research Agency (ANR-EPIBACTIN). Wenyang Dong is part of the Pasteur - Paris University (PPU) International PhD Program, a project which has received funding from the European Union’s Horizon 2020 research and innovation programme under the Marie Sklodowska-Curie grant agreement No 665807. Wenyang Dong is supported by the EUR G.E.N.E. (reference #ANR-17-EURE-0013) and is part of the Université de Paris IdEx #ANR-18-IDEX-0001 funded by the French Government through its “Investments for the Future” program. We would like to thank Dr. Patrick Trieu-Cuot, Dr. Thomas Kohler, Dr. Mustapha Si-Tahar, Dr. Emmanuelle Varon and Dr. Birgitta Henriques for providing the strains used in this study.

## Author Contributions

Conceptualization, W.D. and M.A.H.; Methodology, W.D. and M.A.H; Investigation, W.D., O.R., C.C., M.C., and M.E.; Writing – Original Draft, W.D. and M.A.H.; Writing –Review & Editing, W.D. O.R., C.C., M.C., M.E., and M.A.H.; Funding Acquisition, M.A.H.; Supervision, M.A.H.; Project Administration, M.A.H.

## Declaration of Interests

The authors declare no competing interests.

## Materials and methods

### Antibodies

Antibodies used in this study: anti-actin (Sigma, A5441), anti-H3S10ph (Millipore, MC463), anti-H3 (AbCam, ab1791), anti-H3T3ph (AbCam, ab78531), anti-H3T6ph (AbCam, ab222768), anti-H3T11ph (AbCam, ab5168), anti-H3S28ph (AbCam, ab5169), anti-PP1 that specifically recognizes PP1a (Santa Cruz, sc-7482), anti-PP1b that cross-reacts with both PP1b isoform and PP1c isoform proteins (Millipore, 07-1217), anti-phospho-PP1a (Thr320) antibody that recognizes all PP1 isoforms (Cell Signaling, 2581s) (20), goat anti-rabbit AF647 (Invitrogen, A27040), anti-AKT (Cell Signaling, 2920S), anti-phospho-AKT-S473 (Cell Signaling, 9271S), anti-phospho-(Ser/Thr) Phe (Cell Signaling, 9631S), anti-CD326 (Miltenyi Biotec, REA977), anti-CD31 (Miltenyi Biotec, REA784), anti-CD45 (Miltenyi Biotec, REA 737).

### Chemical and biological and reagents

For experiments involving chemical inhibitors, cells were pretreated for 3h before infection with Tautomycetin (0.8 μM, Tocris Bioscience, 119757-73-2) or Okadaic acid (0.1 μM, Sigma, N0636). Hydrogen peroxide (Sigma, 7722-84-1) was added to the cells for 1h at indicated concentrations. Purified PLY and LLO were obtained as described previously (8). Cells were treated with 6 nM LLO or 6 nM PLY for 20 or 30 min.

### Construction of S. pneumoniae mutants

All primers used in construction of *S. pneumoniae* mutants are listed in Table S1. Mutants of *S. pneumoniae* were generated by transforming donor DNA into pneumococci using competence stimulating peptide (CSP) as described previously (38). To create a deletion allele with selection marker, antibiotic-resistance cassettes were fused with the upstream and downstream flanking regions of target genes by overlapping PCR. Specifically, *ply* gene was replaced by with *erm* cassette in Δ*ply* mutant; *lytA* gene was replaced by with *erm* cassette in Δ*lytA* mutant; in Δ*spxB* mutant, *spxB* gene was replaced by *kan*+*sacB* (KS) cassette, which is a cassette amplified from Sweet Janus cassette (39). The KS cassette permits allelic replacement or marker-free knock in through sequential positive and negative selection. Therefore, we constructed *in situ* complement strains using KS cassette. Briefly, using kanamycin-resistance positive selection, targeted sequence was replaced by KS cassette in chromosome. Using sucrose sensitivity negative selection, KS cassette was further replaced by either wild-type genes or genes carrying point mutation. Point mutation was also created by overlapping PCR.

### Bacterial culture and cell infections

All *S. pneumoniae* strains used in this study are listed in Table S2. *S. pneumoniae* strains were grown in Todd Hewitt broth (Bacto, BD, USA) supplemented with 50 mm HEPES at 37°C with 5% CO2 until the optical density at 600 nm = 0.6. *L. monocytogenes* strains were grown in brain-heart infusion medium (Difco, BD, USA) at 37°C with 5% CO2 until the optical density at 600 nm = 1. Bacteria were washed twice with PBS and diluted in serum-low cell culture medium. For *S. pneumoniae* strains, a multiplicity of infection (MOI) of 50:1 for R6 and a MOI of 25:1 for TIGR4 were used unless otherwise indicated. After 3h of *S. pneumoniae* infection, cells were washed with PBS for 3 times, and then either collected for future process, or cultured in medium with penicillin (10 μg /ml) and gentamicin (100 μg /ml) for later time points. For *L. monocytogenes* strains, which are described previously (8,40), a multiplicity of infection (MOI) of 50:1 was used. After 1h of Listeria infection, cells were washed with PBS 3 times, and cultured in medium with 10 μg/ml gentamycin to carry out Listeria infection for late time points.

The adherence and intracellular assays of *S. pneumoniae* were performed using A549 cells. A total of 5 × 10^6 R6 pneumococci were added to 24-well tissue culture plates containing 1 × 10^5 cells each well. After 3h infection, cells were washed 3 times with PBS to remove the unattached bacteria. To determine the amount of adherent pneumococci, PBS washed cells were lysed using sterile ddH2O. Lysates and its serial dilutions were plated on Columbia blood agar plates (43059, BIOMERIEUX) overnight at 37°C with 5% CO2, and the colony forming units (CFUs) were counted as bacteria adherence. To determine the amount of intercellular pneumococci, PBS washed cells were cultured in in medium with penicillin (10 μg /ml) and gentamicin (100 μg /ml) to kill extracellular bacteria. Sterile ddH2O was added at 4h time point after infection (namely 1h after PBS washing) or 6h time point after infection. Lysates and its serial dilutions were plated on Columbia blood agar plates overnight at 37°C with 5% CO2, and the colony forming units (CFUs) were counted as intercellular bacteria.

### Cell Culture

The human alveolar epithelial cell line A549 (ATCC CCL-185) cells were cultured in F-12K culture medium supplemented with 10% fetal calf serum (FCS) and 1% glutamine. The human bronchial epithelial cell line BEAS-2B (ATCC CRL-9609) cells were cultured in DMEM culture medium supplemented with 10% FCS and 1% glutamine. The human cervical carcinoma epithelial cell line HeLa (ATCC CCL-2) cells and human Colon Carcinoma cell line CaCO2 (ATCC HTB-37) cells were cultured in MEM culture medium supplemented with 1% glutamine, 1 mM sodium pyruvate (GIBCO), 0.1 mM nonessential amino acid solution (GIBCO), and 10% (HeLa) or 20% (CaCO2) FCS. Cells were seeded in 6-well or 24-well plates 2 days before infection. When cells were grown to semi-confluence (24h time point), they were serum-starved (0.25% FCS) for 24h before use in experiments.

### In vivo infections

All protocols for animal experiments were reviewed and approved by the CETEA (Comité d’Ethique pour l’Expérimentation Animale - Ethics Committee for Animal Experimentation) of the Institut Pasteur under approval number Dap170005 and were performed in accordance with national laws and institutional guidelines for animal care and use. Wildtype C57BL/6 female 8-9 week old mice purchased from Janvier Labs (France), *In vivo* infections were performed by intranasal injection of 5×10^5 bacteria per mouse Lungs were collected after 20h from infection and digested using the Lung Dissociaton Kit according to the manufacturer’s instructions (Miltenyi Biotec) plus added Dispase II at 0,1U/ml (Roche). Single cell suspensions were stained to identify lung epithelial cells (CD45-CD31-CD326+). Cell were further permeabilized with Transcription Factor Staining Buffer Set (eBioscience) and stained for H3S10ph followed by secondary. Data was acquired on a MACSQuant cytometer (Milteny Biotec).

### Cell synchronization and cell cycle analysis

Cells were synchronized by a thymidine block as described (41). For cell cycle distribution, cells were detached in PBS and fixed in 70% ethanol for 1h at −20°C. Cells were washed and re-suspended in PBS containing 10 μg/ml of propidium iodide and 100 μg/ml of RNase (DNase free).

### FACS Analyses

Cells infected with GFP-expressing *L. monocytogenes*, cells stained with propidium iodide, and cells from lung of infected mice and stained with antibodies, were analyzed on a FACSCalibur. Data was analyzed using the FlowJo software.

### Transfection of siRNA

Lipofectamine 2000 (Invitrogen, 11-668-019) was used to introduce interference RNA into A549 cells. Pan PP1 siRNA (Santa Cruz, sc-43545) was transiently transfected (25 nM final concentration in well) to knockdown all isoforms of PP1. Scramble siRNA (On-TARGETplus SMARTpool) was transiently transfected as control. Cells were assayed 48h after siRNA transfection.

### Immunofluorescence

Cells were grown on glass cover slides. After infection, cells were washed 3 times with PBS and fixed in 4% paraformaldehyde for 10 min at room temperature. After 3 washes, cells were permeabilized and blocked with 3% BSA with 0.5% Tween20 for 45 min. Immunostaining was performed with primary antibodies in 3% BSA+ 0.5% Tween20 for 90 min, and then Alexa Fluor 488 or 647 conjugated anti-immunoglobulin G (IgG) secondary antibodies in 3% BSA+ 0.5% Tween20 for 45 min.

### Western blot analyses

For Western blot analysis, cells were lysed using 2× laemmli buffer (4% SDS, 20% glycerol, 200mM DTT, 0.01% bromphenol blue and 0.1 M Tris HCl, pH 6.8). Samples were sonicated for 5s, boiled for 10 min, and then subjected to SDS-PAGE. Proteins were transferred to membrane under 2.5A/25V condition for 7min using a semidry transfer system (Trans-Blot Turbo, BIO-RAD). Transferred membranes were first blocked by TBS-Tween20 (0.1%) with 5% milk, and then incubated with primary antibodies overnight at 4°C. Membranes were washed with TBS-Tween20 (0.1%) and incubated with secondary peroxidase-conjugated anti-immunoglobulin G (IgG) antibodies for 1h at room temperature. After washing with TBS-Tween20 (0.1%), the immunoreactive bands were visualized using ECL substrate (Clarity Western ECL substrate, BIO-RAD) and imaged with a Western Blot detection system (ChemiDoc Imaging Systems, BIO-RAD). Quantification of Western blots was performed using Image Lab (BIO-RAD).

### RNA extraction and Quantitative PCR analyses

The mRNA was extracted from cells using an RNeasy kit (Qiagen, 74104). DNase treatment was performed using a DNase set (Qiagen, 79254). cDNA was then synthesized using 1 μg input RNA by BIO-RAD iScript gDNA Clear cDNA Synthesis Kit (1725035). Real-time PCR was performed using using the SYBR Green kit (iTaq Universal SYBR Green Supermix, BIO-RAD, 1725124) on a BIO-RAD CFX384. Data was obtained using BIO-RAD CFX manager. Relative gene expression analysis was performed using the 2^-ΔCT method as described (42). All primers used in quantitative PCR analyses are listed in Table S3.

