## Supplementary figures and images for "*Streptococcus pneumoniae* infection promotes histone H3 dephosphorylation by modulating host PP1 phosphatase"

### Supplemental Figure S1

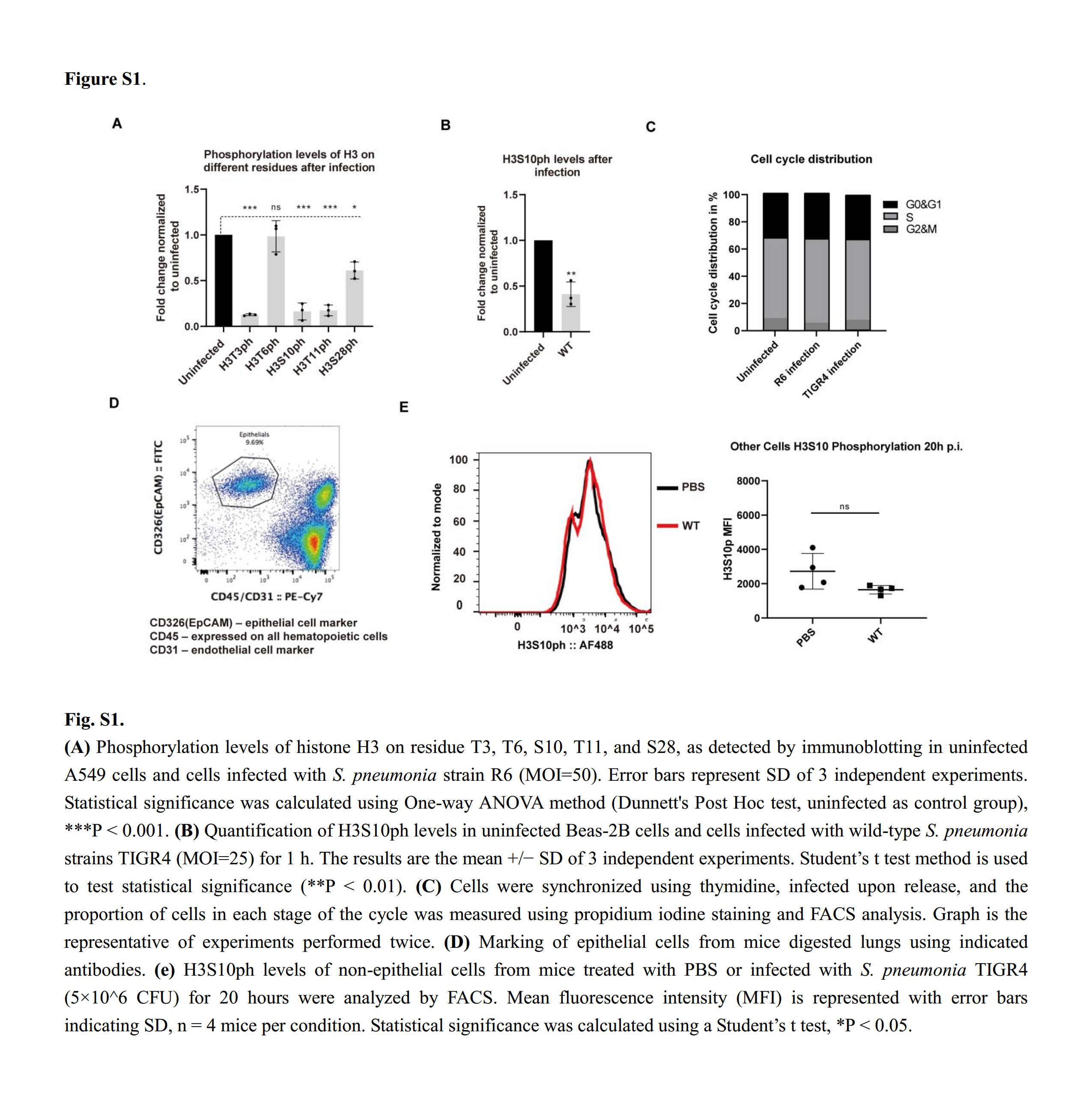

### Supplemental Figure S2

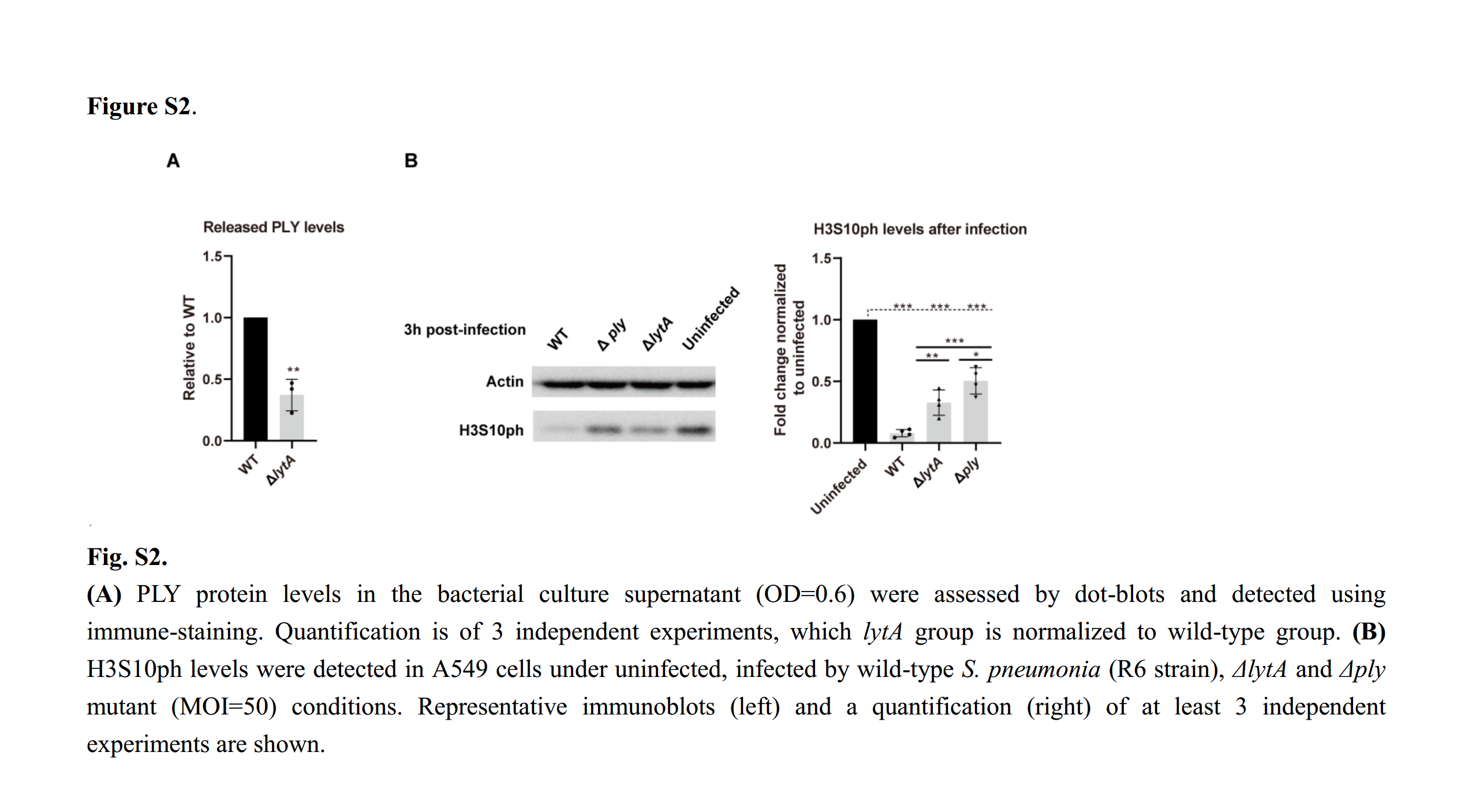

### Supplemental Figure S3

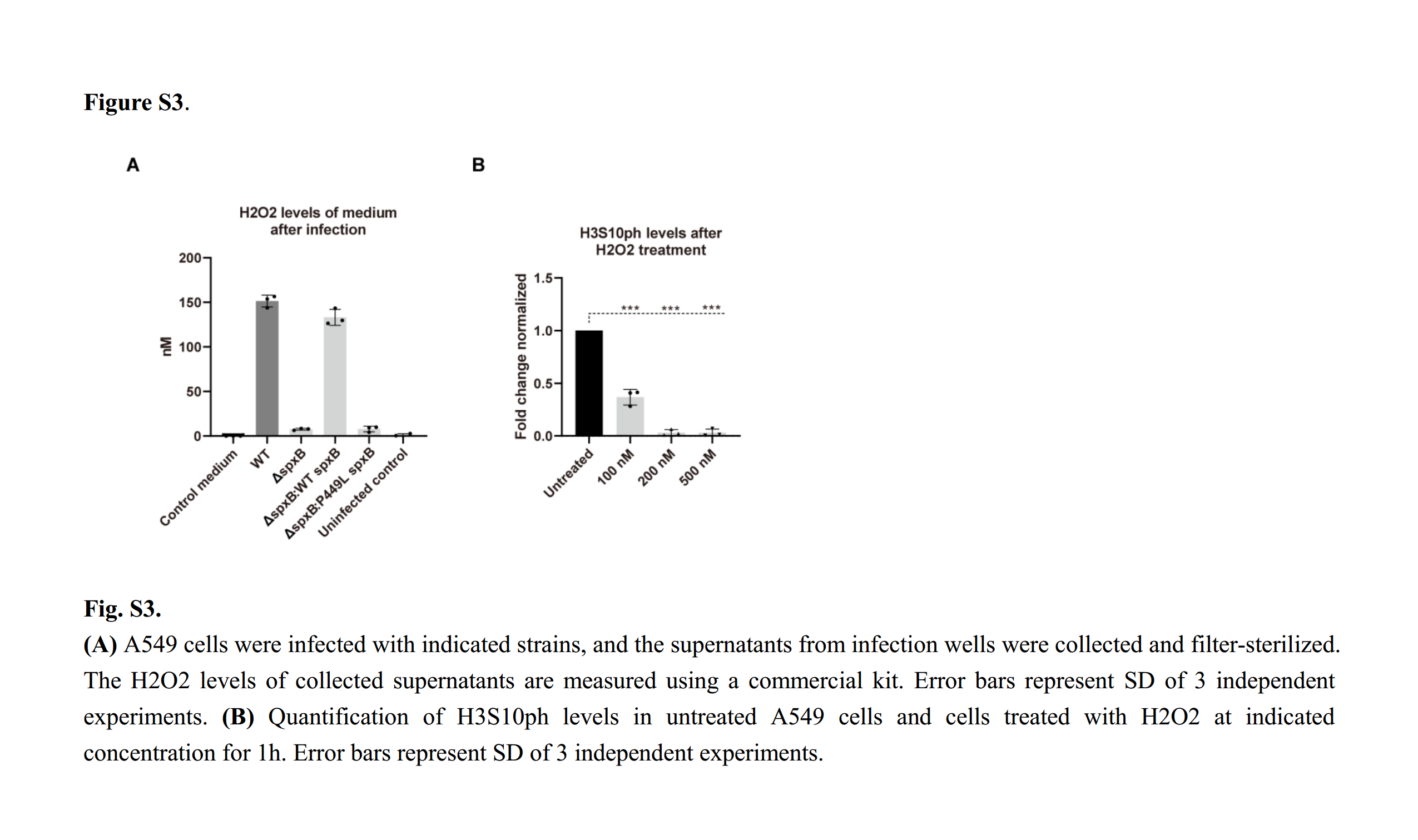

### Supplemental Figure S4

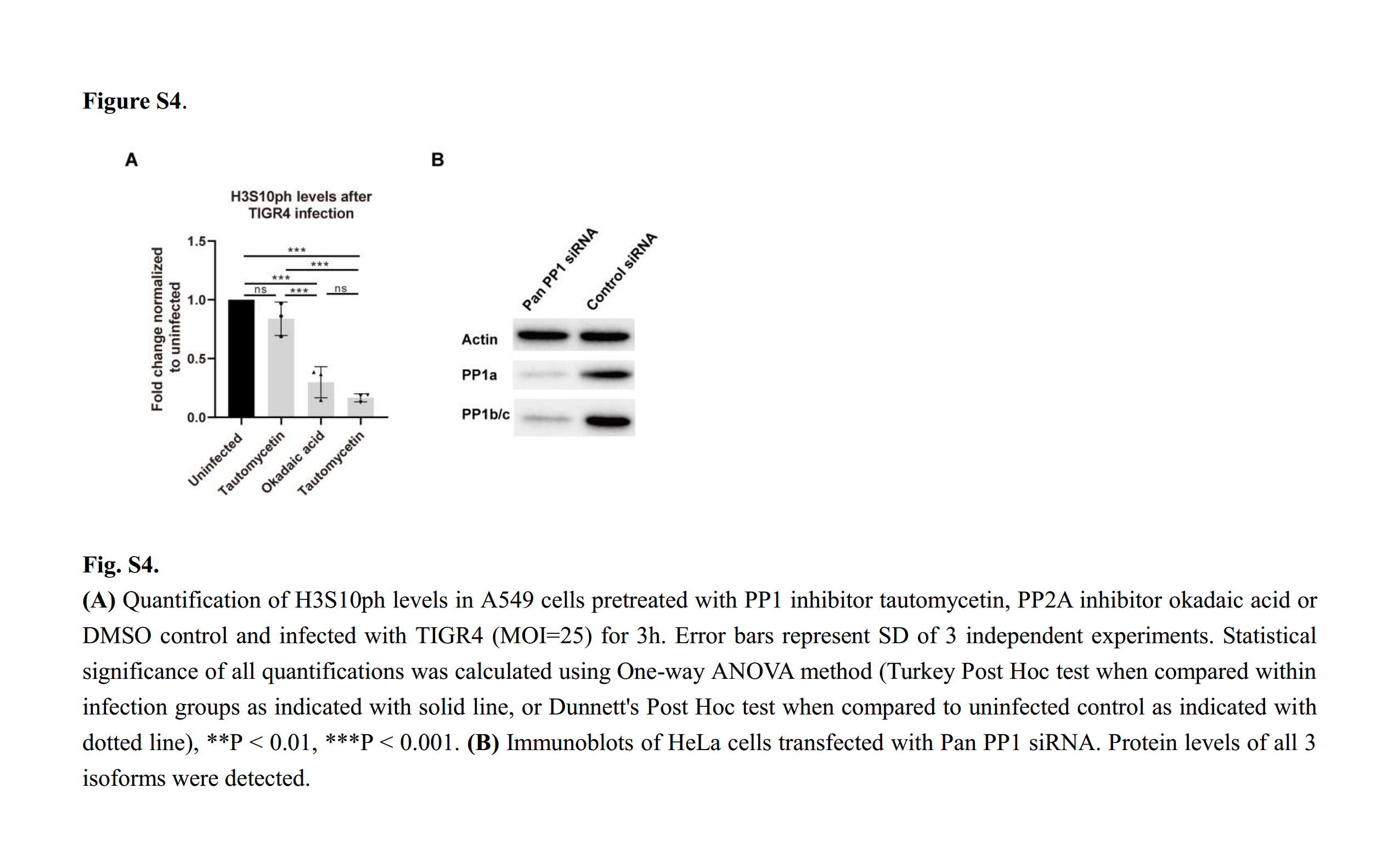

### Supplemental Figure S5

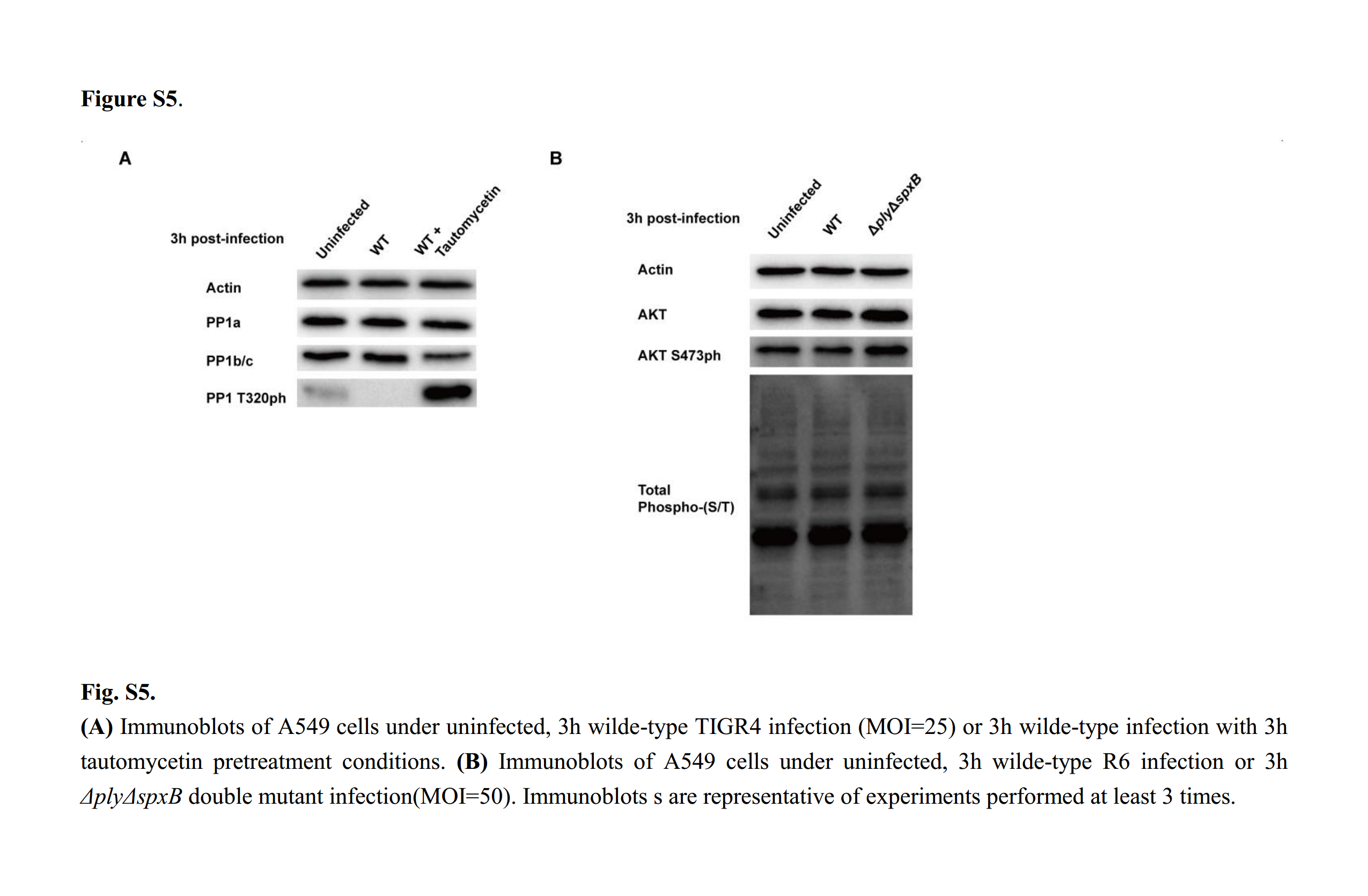
