## Supplemental Tables for "*Streptococcus pneumoniae* infection promotes histone H3 dephosphorylation by modulating host PP1 phosphatase"

Table S1

| Primers | Sequences | Amplification |
| --- | --- | --- |
| PLY-UP- F | TTGGCGACAAGCATTTTGTA | Upstream region of ply |
| PLY-UP-ERM-R | TCTTAATTACAAATTTTTCCACTACGAGAAGTG | Upstream region of ply |
| PLY-ERM-F | CACTTCTCGTAGTGGAAAAATTTGTAATTAAGAAGGAG | Cassette ERM in ply mutant |
| PLY-ERM-R | CGAACATTCCCTTTTCCAAATTTAAAAAAGCG | Cassette ERM in ply mutant |
| PLY-DOWN-ERM-F | GCTTTTTTAAATTTGGCTTTAAAAGGGAATG | Downstream region of ply |
| PLY-DOWN-R | GCGACAAAAACAATCATACTGC | Downstream region of ply |
| PLY-KS-F | CTGGAGCACTGGATAATGCTGAAAACTCCTT | Cassette KS in ply mutant |
| PLY-KS-R | TCCTCCCGTTAGTACTAAAACAATTCATCCAG | Cassette KS in ply mutant |
| PLY-UP-SK-R | AGCATTATCCAGTGCTCCAGGATAGAGGCGACTG | Upstream region of ply |
| PLY-DOWN-SK-F | GTTTTAGTACTAACGGGAGGAAATAATTCTATG | Downstream region of ply |
| PLY-436A-F | GTACGCGCCCATTCCCAGGCAAGCCC | Single point mutation in ply |
| PLY-436A-R | TGCCTGGGAATGGGCGCGTACGGTT | Single point mutation in ply |
| LytA-UP-F | GCCAGTCCAGCTTTGGTTTCCTT | Upstream region of lytA |
| LytA-UP-R | TCTTAATTACAAATTTTTCCAGAACCAGAAACTCC | Upstream region of lytA |
| LytA-ERM-F | GGAGTTTCTGGTTCTGGAAAAATTTGTAATTAAGAAGGA | Cassette ERM in lytA mutant |
| LytA-ERM-R | CTGATTTGAAAGACATTCCCCAAATTTAAAAAAGCG | Cassette ERM in lytA mutant |
| LytA-DOWN-F | CGCTTTTTTAAATTTGGGGAATGTCTTTCAAATCAG | Downstream region of lytA |
| LytA-DOWN-R | GTCTGAATTGACTGCAATGAGACTGAAC | Downstream region of lytA |
| SpxB-UP-F | CGTGGTCTCCGAACAGTCATGC | Upstream region of spxB |
| SpxB -UP-R | ATCAAACGGAATAATGATAACTCTCCTTCAATTT | Upstream region of spxB |
| SpxB -KS-F | TTATCATTATTCCGTTTGATTTTTAATGGATA | Cassette KS in spxB mutant |
| SpxB -KS-R | GTACGAGTTCGTACTAAAACAATTCATCCAG | Cassette KS in spxB mutant |
| SpxB -DOWN-F | GTTTTAGTACGAACTCGTACCATTCCGTCTC | Downstream region of spxB |
| SpxB -DOWN-R | GAACCCCATTGAACAAGTGTG | Downstream region of spxB |
| SpxB –P449L-F | TGGAGCATTCAACATGTGCTACCTAGA | Single point mutation in spxB |
| SpxB –P449L-R | TCTAGGTAGCACATGTTGAATGCTCCA | Single point mutation in spxB |

Table S2

| Strains/serotypes | MLST | Source |
| --- | --- | --- |
| R6 (uncapsulated) | ST595 | Laboratory strain, gifted by Dr. Patrick Trieu-Cuot |
| TIGR4 (serotype 4) | ST205 | Laboratory strain, gifted by Dr. Thomas Kohler |
| Serotype 1 | ST304 | Invasive clinical strain, gifted by Dr. Mustapha Si-Tahar |
| Serotype 1 | ST306 | Invasive clinical strain, gifted by Dr. Emmanuelle Varon |
| Serotype 3 | ST180 | Invasive clinical strain, gifted by Dr. Emmanuelle Varon |
| Serotype 3 | ST260 | Invasive clinical strain, gifted by Dr. Emmanuelle Varon |
| Serotype 7F | ST191 | Invasive clinical strain, gifted by Dr. Emmanuelle Varon |
| Serotype 6B | ST90 | Carriage clinical strain, gifted by Dr. Emmanuelle Varon |
| BHN78 (serotype 14) | ST124 | Carriage clinical strain, gifted by Dr. Birgitta Henriques |
| Serotype 19A | ST276 | Carriage clinical strain, gifted by Dr. Emmanuelle Varon |
| BHN100 (serotype 19F) | ST162 | Carriage clinical strain, gifted by Dr. Birgitta Henriques |

Table S3

| RT-qPCR primers | Sequences |
| --- | --- |
| GAPDH-F | ACATCGCTCAGACACCATG |
| GAPDH-R | TGTAGTTGAGGTCAATGAAGGG |
| HPRT1-F | TGCTGAGGATTTGGAAAGGG |
| HPRT1-R | ACAGAGGGCTACAATGTGATG |
| CCL2-F | TGTCCCAAAGAAGCTGTGATC |
| CCL2-R | ATTCTTGGGTTGTGGAGTGAG |
| CCL4-F | CTGTGCTGATCCCAGTGAATC |
| CCL4-R | TCAGTTCAGTTCCAGGTCATACA |
| CCL5-F | CCAGCAGTCGTCTTTGTCAC |
| CCL5-R | CTCTGGGTTGGCACACACTT |
| CSF2-F | TCCTGAACCTGAGTAGAGACAC |
| CSF2-R | TGCTGCTTGTAGTGGCTGG |
| IL1A-F | AGATGCCTGAGATACCCAAAACC |
| IL1A-R | CCAAGCACACCCAGTAGTCT |
| CXCL1-F | AACCGAAGTCATAGCCACAC |
| CXCL1-R | CCTCCCTTCTGGTCAGTTG |
| TNF-F | ACTTTGGAGTGATCGGCC |
| TNF-R | GCTTGAGGGTTTGCTACAAC |
| DEFA1-F | GGACATCCCAGAAGTGGTTG |
| DEFA1-R | GTAGATGCAGGTTCCATAGCG |
| CAMP-F | TGTGCTTCGTGCTATAGATGG |
| CAMP-R | GCACACTGTCTCCTTCACTG |
| DEFB4-F | CCATGAGGGTCTTGTATCTCC |
| DEFB4-R | AGGGCAAAAGACTGGATGAC |
| DEFA4-F | GCCAGAAGACCAGGACATATC |
| DEFA4-R | ATGAGGCAGTTCCCAACAC |
| DEFB1-F | GCCATGAGAACTTCCTACCTTC |
| DEFB1-R | CAGAATAGAGACATTGCCCTCC |
| CXCL2-F | AACCGAAGTCATAGCCACAC |
| CXCL2-R | CTTCTGGTCAGTTGGATTTGC |
| CXCL5-F | TCTGCAAGTGTTCGCCATAG |
| CXCL5-R | CAGTTTTCCTTGTTTCCACCG |
| IL6-F | CCACTCACCTCTTCAGAACG |
| IL6-R | CATCTTTGGAAGGTTCAGGTTG |
| IL17C-F | GAGGTGTTGGAGGCAGAC |
| IL17C-R | CAGCTTCTGTGGATAGCGG |
| IL25-F | GTGAAGATGGACCCCTCAAC |
| IL25-R | CTGTAGGCTGACGCAGTG |
| IL15-F | ACCGTGGCTTTGAGTAATGAG |
| IL15-R | AAGCCCTGCACTGAAACA |
| IL33 | AGTCTCAACACCCCTCAAATG |
| IL33 | CTTTTGTAGGACTCAGGGTTACC |
| TNFRSF1A -F | TGCCAGGAGAAACAGAACAC |
| TNFRSF1A –R | TCCTCAGTGCCCTTAACATTC |
| IL1R1 | GATGAAGATGACCCAGTGCTAG |
| IL1R1 | TGGCAAAACAGGTAAATGGATG |
| TLR1-F | CCCGGAAAGTTATAGAGGAACC |
| TLR1-R | CAGATCCAAGTAGCTGCAGAG |
| TLR2-F | TGGTAGTTGTGGGTTGAAGC |
| TLR2-R | GACAGAGAAGCCTGATTGGAG |
| TLR4-F | TGCGTGAGACCAGAAAGC |
| TLR4-R | TTAAAGCTCAGGTCCAGGTTC |
| TLR5-F | GCTAGGACAACGAGGATCATG |
| TLR5-R | GAGGTTGCAGAAACGATAAAAGG |
| IL17RE-F | AATTCCTTCTGCCCTGTCTG |
| IL17RE-R | TCCAGAAGTCCGAGCCATAG |
